# Cyclic mechanical stresses alter erythrocyte membrane composition and microstructure and trigger macrophage phagocytosis

**DOI:** 10.1101/2021.09.10.459518

**Authors:** Antoni Garcia-Herreros, Yi-Ting Yeh, Zhangli Peng, Juan C. del Álamo

**Affiliations:** Department of Mechanical and Aerospace Engineering, University of California, San Diego, La Jolla, CA, USA; Department of Bioengineering, University of California, San Diego, La Jolla, CA, USA; Institute of Engineering in Medicine, University of California, San Diego, La Jolla, CA, USA; Department of Bioengineering, University of Illinois at Chicago, Chicago, IL, USA; Department of Mechanical Engineering, University of Washington, Seattle, WA, USA; Center for Cardiovascular Biology, University of Washington, Seattle, WA, USA

**Keywords:** Red blood cells, cell aging, Inter endothelial slits, Vesiculation, Microfluidics, Macrophage phagocytosis

## Abstract

Red blood cells (RBCs) are cleared from the circulation when they become damaged or display aging signals targeted by macrophages. This process occurs mainly in the spleen, where blood flows through submicrometric constrictions called inter-endothelial slits (IES), subjecting RBCs to large-amplitude deformations. In this work, we circulated RBCs through microfluidic devices containing microchannels that replicate the IES. The cyclic mechanical stresses experienced by the cells affected their biophysical properties and molecular composition, accelerating cell aging. Specifically, RBCs quickly transitioned to a more spherical, less deformable phenotype that hindered microchannel passage, causing hemolysis. This transition was associated with the release of membrane vesicles, which self-extinguished as the spacing between membrane-cytoskeleton linkers became tighter. Proteomics analysis of the mechanically aged RBCs revealed significant losses of essential proteins involved in antioxidant protection, gas transport, and cell metabolism. Finally, we show that these changes made mechanically aged RBCs more susceptible to macrophage phagocytosis. These results provide a comprehensive model to explain how physical stress induces RBC clearance in the spleen.

## Introduction

Circulating red blood cells (RBCs) deliver oxygen from the lungs to all organs in the body. Human RBCs are anucleate and discoidal in shape, with an 8-µm diameter and a thickness ranging between 1 and 2.2 µm^1^. Throughout their 120-day lifetime, RBCs undergo large-amplitude deformations in the systemic and pulmonary microcirculations. However, the most severe deformations occur as blood transits the spleen, where RBCs squeeze through 0.5-2.5 µm-gaps between specialized endothelial cell fibers called inter-endothelial slits (IES)^2^. Old and diseased RBCs are retained during splenic transit and eventually removed by red pulp macrophages. The interplay between RBC deformation, RBC aging, and splenic clearance is crucial in hematologic disorders and the preservation of blood stored for transfusion.

As RBCs age, their shape transitions from biconcave-discoidal to serrated or spherical, and their intracellular density increases^3, 4^. These changes, often attributed to membrane loss, are associated with decreased deformability, leading to failure to squeeze through the IES^5^. RBC aging hallmarks are accentuated in hemolytic diseases like iron deficiency anemias and malaria^6^, which cause life-threatening symptoms and multiple complications. Additionally, aged RBCs exhibit significant biochemical changes affecting cell volume, deformability, and metabolic activity, such as ATP depletion^7^ and increased cytoskeleton oxidation^8^. Oxidation, which is also linked to RBC aging during storage^9^, promotes the aggregation of band3^10^, a membrane protein required for CO_2_ transport that regulates RBC stiffness by anchoring the lipid bilayer to the cytoskeleton.

How macrophages identify and remove RBCs from the circulation is still poorly understood. This specific macrophage attack is associated with the presence of distinctive age-related signals in the RBC membrane. Previous studies reported an increase in phosphatidylserine (PS) in the outer leaflet of the membrane as RBCs age^11^. PS is a phospholipid usually located in the inner leaflet of the membrane; however, when exposed in the outer leaflet, it acts as a phagocytosis signal^12^. Additionally, the expression of CD47, a membrane protein that inhibits macrophage attacks, declines as RBCs age^13^. Finally, band3 clustering promoted by age-induced oxidation facilitates autoantibody binding and could mediate RBC recognition by macrophages^14^.

Increased intracellular density is the primary experimental marker of RBC aging because it correlates with other hallmarks of RBC functional decline. Recent data suggest that RBCs exposed to cyclic stretch exhibit significant changes in shape and mechanical properties (i.e., mechanical fatigue)^15,16^. This evidence led us to hypothesize that RBC deformations cause biophysical and biochemical changes equivalent to aging, which can be accelerated significantly in vitro. The main goals of the present study were to test these hypotheses and to investigate the underlying mechanisms for mechanically induced RBC aging. To this end, we fabricated PDMS-based microfluidic devices to model the passage of RBCs through IES in the spleen. We analyzed the biophysical and biochemical changes undergone by RBCs as a function of the number of repetitive passages through these microchannels, finding significant metabolic, proteomic, and biomechanical alterations. Specifically, we demonstrated that RBCs undergo cellular vesiculation and cell rounding accompanied by key biological hallmarks of RBC aging like decreased intracellular ATP and enhanced oxidation, all associated with a higher susceptibility to macrophagic attack. In addition to providing insights into the biomechanical underpinnings of RBC aging, this work offers a proof of principle of using microfluidic devices to systematically and reproducibly accelerate RBC aging *in vitro*.

## Results

### Microfluidic devices mimic crossing of splenic inter-endothelial slits by RBCs

We developed microfluidic devices to model the crossing of RBCs through splenic inter-endothelial slits (IES) using a two-step photolithography process (see Materials and Methods and Fig 1A-B). The devices test section contains an array with 100 microchannels of tunable width (Fig 1 C). Microchannels with widths < 1 μm were achieved as measured by scanning electron microscopy (Supp Fig 1A).

**Figure 1.**
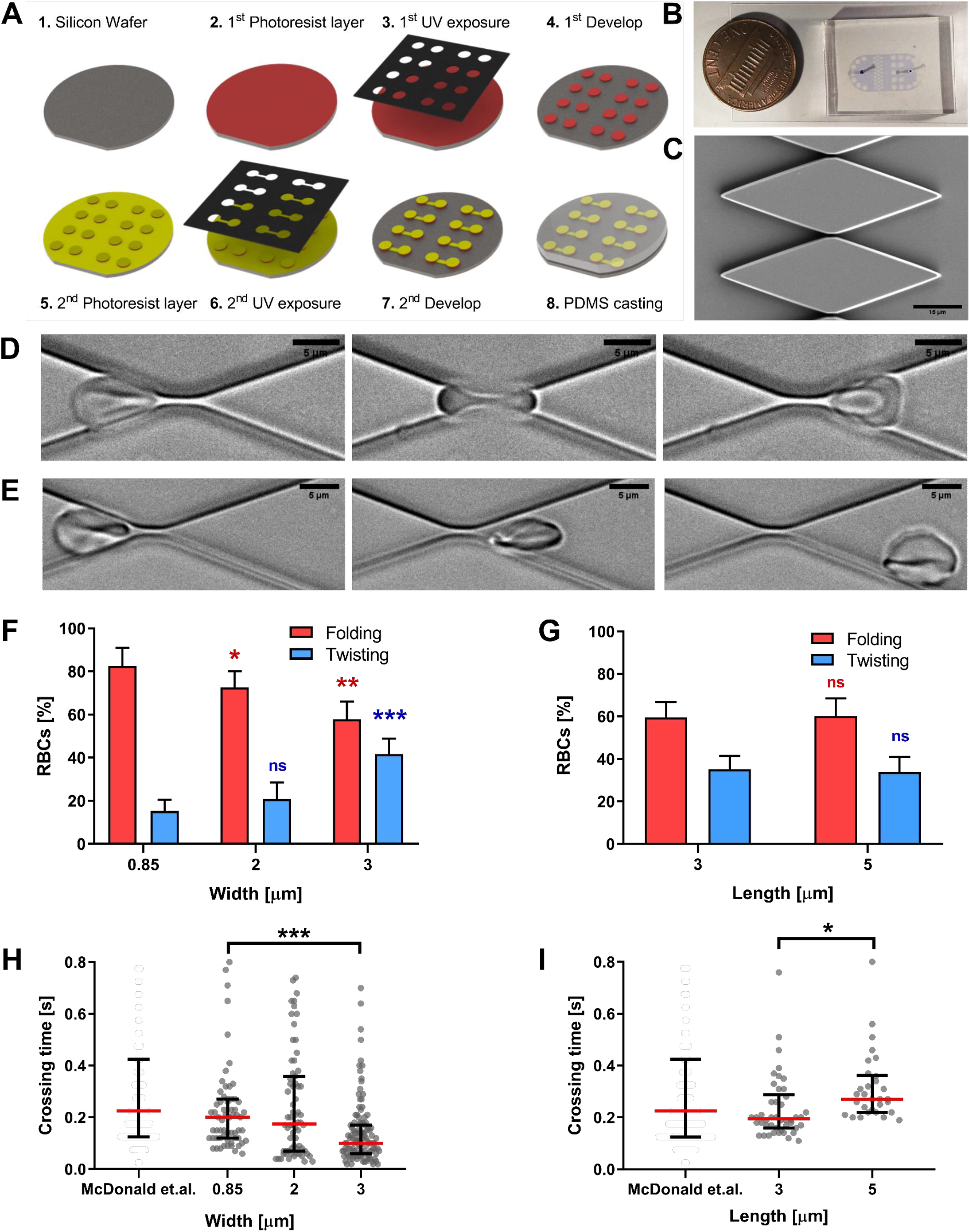
Design of a microfluidic device to replicate the passage of RBCs through splenic inter endothelial slits. **A**, Scheme of the key steps in the fabrication process. See details in the Materials and Methods section. **B**, Actual device next to 1-cent coin for scale. **C**, Scanning electron microscopy image showing a close-up of the micro-constriction array. **D-E**, Time-lapse sequences of RBC passage reveal two characteristic types of motion: folding (D) and twisting (E). See Suppl. Videos 1 and 2 for a high-time-resolution depiction of these two motions. Bars correspond to 15 μm (C) and 5 μm (D and E). Type of RBCs motion was measured as a function of slit width (**F**) or length (**G**). RBCs crossing times were measured with respect to slit width (**H**) and length (**I**) and compared with in vivo data previously calculated by other authors^20^. Each cell is represented as a circle; error bars represent median and interquartile range. Statistically significant differences, as determined using an unpaired t test with Welch’s correction, are indicated (*, p< 0.05; **, p<0.01, ***, p< 0.001).

The small height of the device’s test section forced the RBCs to orient their discoidal plane parallel to the top and bottom walls, subjecting the cells to large-amplitude deformations when crossing the microchannels. Two types of crossing motion were observed. Most often, the RBCs folded like a “taco tortilla” (Fig 1D and Suppl Video 1), a behavior commonly observed in capillaries^17^. Less frequently, the RBCs twisted to progressively align their discoidal plane parallel to the microchannel sidewalls (Fig 1E and Suppl Video 2). A small percentage of cells failed to cross, got trapped in the microchannel, and eventually burst. By varying microchannel geometry within a range representative of IES *in vivo* observations^18^, we found that the frequency of twisting increased with microchannel width (Fig 1F). On the other hand, microchannel length did not affect the type of RBC crossing motion (Fig 1G).

We determined the RBC crossing time through microchannels, a parameter strongly related to cell deformability^19^, as a function of microchannel width and length (Fig 1H-I). We used this information to choose physiologically realistic microchannel dimensions for our experiments that yielded crossing times consistent with available *in vivo* data^20^.

### Successive crossing of microchannels promotes RBC aging

In humans, RBCs recirculate through the IES in the spleen approximately every 200 minutes leading to 400 crossings over a cell’s half-life^21^. To decouple the effects of mechanical deformation from other time-dependent processes, RBCs were recirculated through our microfluidic device to cross the microchannel array every 10 minutes (Suppl Fig 1B). Overall, we recirculated the cells up to 480 times over 3 days through 3×0.85-µm channels. In the control experiments, RBCs were recirculated through a sham microfluidic device without microchannels.

RBC dynamics were significantly affected as the cumulative number of microchannel crossings (*Nc*) increased. The crossing time grew significantly with *Nc* (Fig. 2A), and so did the percentage of cells undergoing folding vs. twisting (Fig. 2B). The number of cells trapped inside the microchannels (Fig. 2C) and RBC lysis also increased with *Nc* (Fig. 2D). Successive microchannel crossing precipitated RBC aging hallmarks such as oxidation (assessed by RBC methemoglobin content, Fig. 2E) and intracellular ATP decline (Fig. 2F). Altogether, these data demonstrate that cyclic large-amplitude loading promotes an aged phenotype in RBCs.

**Figure 2.**
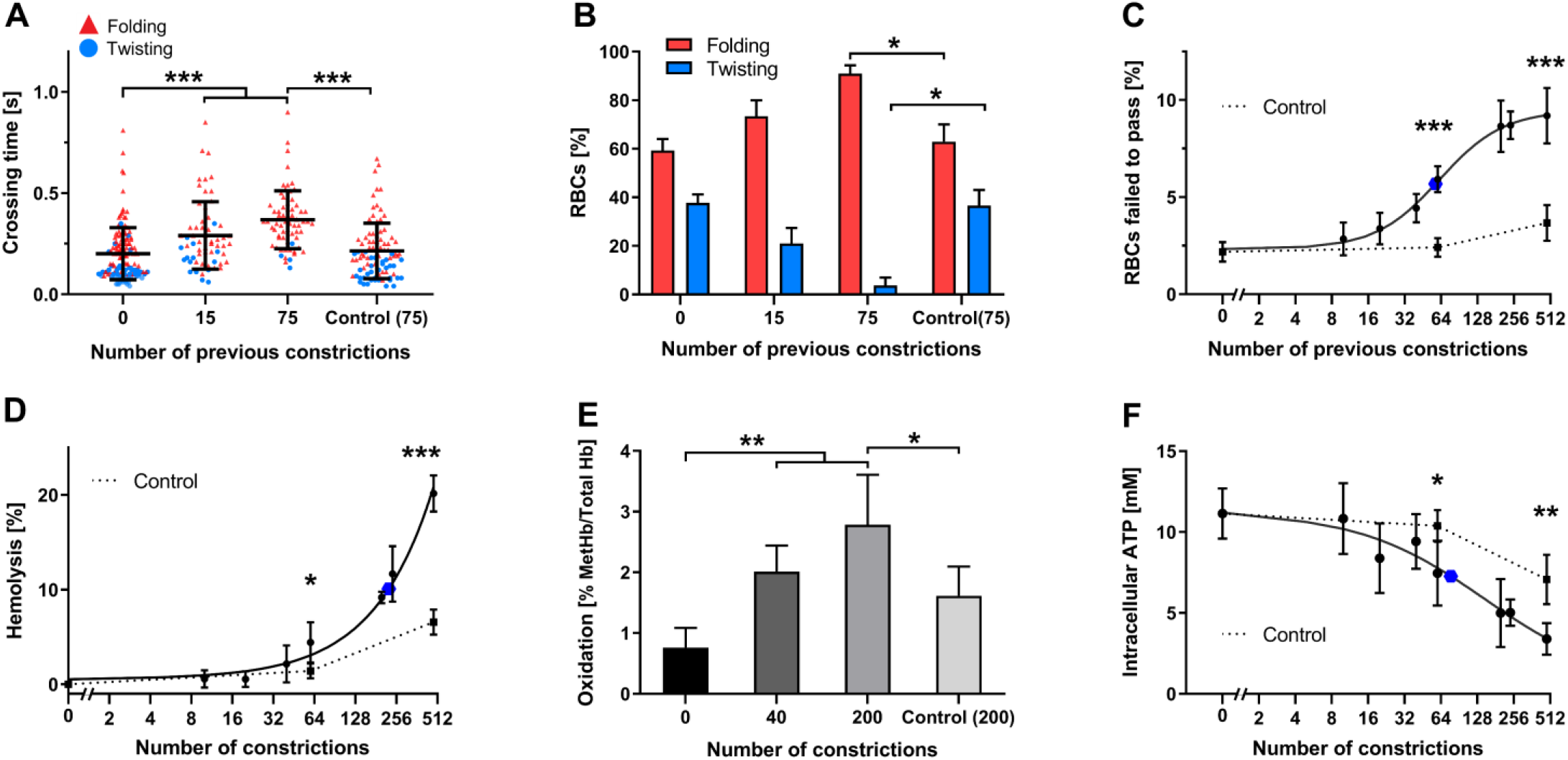
Successive microchannel crossing promotes RBC aging. **A**, Crossing time was measured as indicated in methods in RBCs that had previously gone through 0, 15, or 75 microchannels (0.85µm width and 3µm length) by recirculating in the microfluidic circuit. Each cell is represented as a triangle or circle depending on whether it underwent folding or twisting, respectively. The control corresponds to cells that have gone to the indicated number of previous passages through a sham microfluidic device containing no constrictions. **B**, Percentage of cells displaying folding vs. twisting motion after previous passage through 0, 15, or 75 microchannels. **C**, Percentage of RBCs that failed to pass through a micro-constriction vs. number of previous microchannels they have passed through (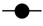,dots, continuous line) and vs. number of previous passages through a mock microfluidic device containing no constrictions (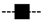, squares, discontinuous line). Data was modeled using the Hill dose-response equation ^55, 56^; **D**, percentage of RBC lysis vs. number of previous microchannels RBCs have passed through (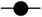, dots, continuous line) and vs. number of previous passages through a mock microfluidic device containing no constrictions (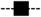 squares, discontinuous line). Data was modeled using a double exponential decay equation. **E**, RBC oxidation vs. number of constrictions the cells have previously passed. **F**, Intracellular ATP concentration vs. number of previous microchannels and vs. number of previous passages through a mock microfluidic device containing no constrictions. Data was modeled using the Hill dose-response equation. In panels A-D, the data come from at least 20 determinations of 3 independent experiments, in E and F, from the evaluation of 5 independent experiments. All datapoints are represented as mean ± SD. In all the graphs the blue dot 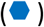 represents the point with the 50% change. Statistically significant differences, as determined using an unpaired t test with Welch’s correction, are indicated (*, p< 0.05; **, p< 0.01; ***, p< 0.001).

### Microchannel crossing results in RBC volume loss and vesiculation

Since RBC aging is associated with changes in cell volume and sphericity^3, 4^, we investigated how the cumulative crossing of microchannels affected these parameters. The sphericity index was defined as *SI* = π^1/3^6^2/3^*V*^2/3^/*S*, where *V* and *S* are cell volume and surface area measured by fluorescence exclusion^22^ (see Materials and Methods, and Suppl Fig. 2A-B). We separated RBCs based on their density by centrifugation to independently study lighter (presumably younger) and denser (older) cells.

Our measurements revealed that cells lost volume and became more spherical as *Nc* increased (Fig. 3A-B). These changes were more significant in lighter cells than in denser ones. In contrast, control cells circulated for an equivalent time through the sham circuit were significantly less affected. After passing through *Nc* ≈ 60 microchannels, the statistical differences between light and dense cells became non-significant, implying that the initially young RBCs became similar to the aged ones. Of note, the RBCs needed to loop in the microfluidic circuit for only 10 hours to achieve the aged phenotype. Microchannel crossing time directly correlated with cell sphericity (Fig. 3C) and a significant proportion of cells with *SI* ≳ 0.8 got stuck inside the microchannels. Consequently, the rate of microchannel RBC retention and hemolysis increased with *Nc* (Suppl Fig. 3A-B). We used membrane flickering spectrometry to assess the RBC membrane (lipid bilayer + membrane skeleton) effective tension and bending modulus. We validated this technique using previously published data for control RBCs^23^ (Suppl Fig. 2C-D). We observed that the bending modulus increased with *Nc* (Fig 3D) while tension was not altered significantly (Fig 3E).

**Figure 3.**
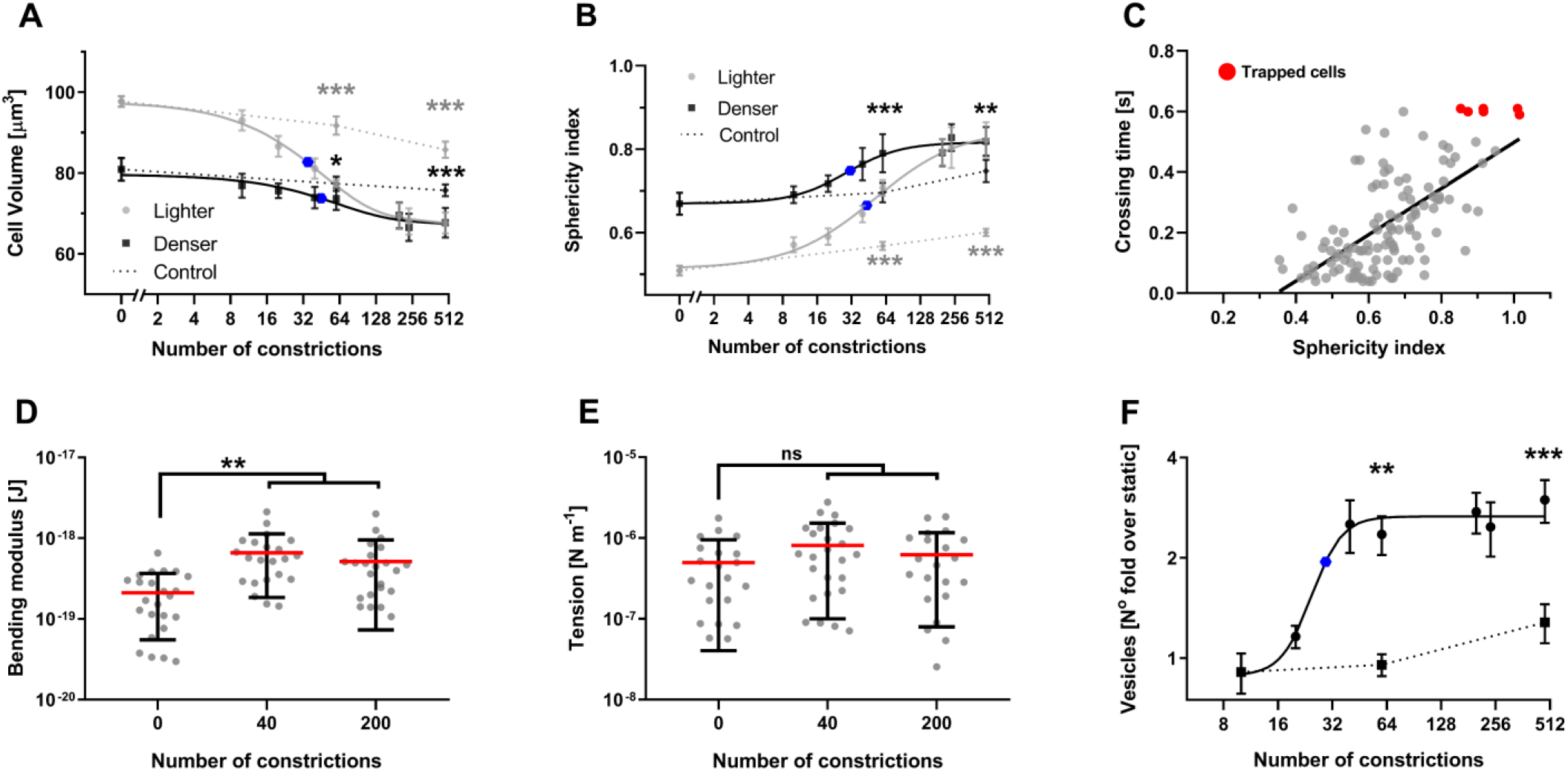
Microchannel crossings result in RBC volume loss associated with vesiculation. **A** and **B**, Two subpopulations of RBCs were separated based on density and circulated multiple times through our microfluidic circuit (constrictions’ dimensions: 0.85µm width and 3µm length). Cell volume (A) and sphericity index (B) were plotted vs. number of microchannels previously passed by RBCs of low-density (grey dots,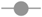) or high-density (black squares,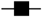) and compared with RBCs recirculated for an equivalent time through a mock microfluidic device containing no constrictions (control, discontinuous line; grey diamonds,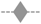, low-density cells; black diamonds,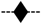, high-density cells). Data was modeled using the Hill dose-response equation. The data are represented as mean and 95% confidence interval of at least 20 determinations of 3 independent experiments. Statistically significant differences, as determined using an unpaired t test with Welch’s correction, are indicated (*, p< 0.05; **, p<0.01; ***, p< 0.001). **C**, Scatter plot of micro-constriction crossing time vs. sphericity index where each point represents a single cell. **D-E**, ‘effective’ bending elastic modulus (D) and cortical tension (E) were assessed in at least 20 RBCs after they passed through the indicated number of constrictions. Each point represents a cell and the error bars indicate mean ± SD (**, p<0.01; Wilcoxon test). **F**, Vesicle release was determined by flow cytometry and represented as fold change relative to static controls. Dotted line corresponds to cells were recirculated for an equivalent time without constrictions. The means ± SEM of 5 independent experiments are shown. Data was modeled using the Hill dose-response equation. In all graphs the blue dot 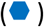 represents the point with the 50% change.

RBC densification and volume loss have been attributed to vesiculation^24^. Therefore, we studied whether cyclic microchannel crossing caused RBC vesiculation. Vesicle generation was assessed by flow cytometry, which was calibrated using microbeads (Suppl Fig 4A-C). Based on this analysis, RBCs generated vesicles with diameters between 0.1 and 0.5 µm, although the release of smaller vesicles undetectable by the flow cytometer cannot be discarded. RBCs subjected to cyclic microchannel crossing released significantly more vesicles than control RBCs kept in stationary conditions (*Nc = 0*) or circulated through the sham device (Fig 3F). The total number of vesicles increased with *Nc* and plateaued around *Nc* ≈ 60, which coincides with the value of *Nc* beyond which RBC volume ceased to decrease (see Fig 3A). Thus, we concluded that volume loss was caused by vesiculation.

### Successive microchannel crossings alter RBC membrane composition and cytoskeletal organization

Our observation that cyclic mechanical loading triggered an aged RBC phenotype motivated a more comprehensive analysis of RBC composition and structure. We used mass spectrometry to analyze the protein content of RBC membranes after 200 microchannel crossings (Fig. 4A, complete list in Suppl Table 1). RBCs kept in stationary conditions (*Nc = 0*) were used as controls. The volcano plot for these proteomics data was asymmetric, with few proteins substantially decreased and many other increased albeit to a lower, not significant extent (Fig. 4B). The list of proteins experiencing statistically significant losses with p-value < 0.01 was analyzed using Metascape^25^ to investigate the most likely affected pathways (Fig. 4C). Protein clustering analysis revealed three affected groups. One group (Log_10_ p-value = −6.2) consisted of proteins involved in cell metabolism like ATP synthase, suggesting a reduction in overall RBC function. This group was tightly linked to a second group (Log_10_ p-value = −5.1) of proteins involved in gas transport and exchange, such as bisphosphoglycerate mutase (BPGM) and carbonic anhydrases. The third group (Log_10_ p-value = −4.6) comprised proteins involved in protection from oxidation such as catalase and peroxiredoxins, congruent with our finding that RBC oxidation increased with *Nc* (Fig. 2E). Although less significant (Log_10_ p-value = −2.8), we also found losses in structural proteins such F-actin, filamin-B, dynactin, or the ankyrin repeat domain. The quantity of other structural proteins like spectrin, band3, band4.1, or ankyrin1 remained unaltered.

**Figure 4.**
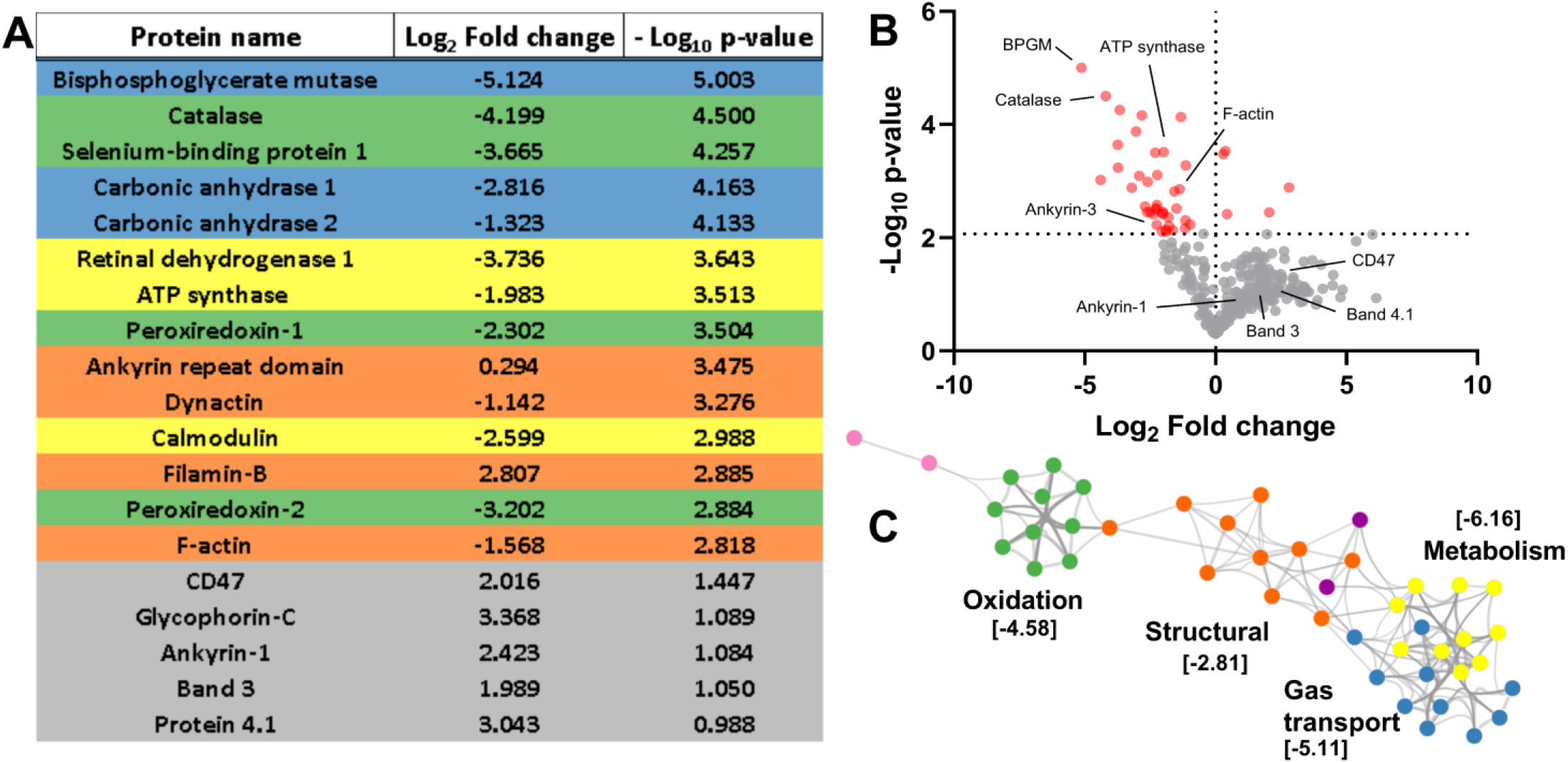
Repetitive microchannel crossings modify RBC protein composition. RBCs were subjected to 200 microchannel crossings (0.85µm width and 3µm length), membranes were isolated, and proteins were extracted and analyzed by mass spectrometry. The differences in protein content between *Nc=200* RBCs and static controls are presented. Values correspond to mean of three independent analysis. **A**, a list of proteins significantly modified (log fold change < −2 or > 2 and a P-value >0.05) and proteins important in membrane cytoskeleton connectivity were classified by its main function. Proteins were classified based on its function, gas transport (blue), metabolism (yellow), structural (orange) and protection from oxidation (green). B, Volcano plot of all the proteins measured. Statistically significant changes were obtained by performing the Benjamini-Hochberg correction method. **C**, Protein clustering analysis revealed that deminished proteins were mostly localized in two clusters: oxidation, gas transport and metabolism. Statistical differences in **C** are indicated as [Log_10_ p-value].

The proteins connecting the RBC lipid bilayer and cytoskeleton, e.g., ankyrin1 or band4.1, form a network with a relatively regular hexagonal pattern^1, 19^. Since successive microchannel crossings did not significantly affect the amounts of these proteins, we hypothesized that their inter-protein spacing in the network shortens as RBC vesiculation reduces the available plasma membrane (Fig 5A). Using stimulated emission depletion microscopy (STED) and image correlation analysis (Fig 5B), we quantified the inter-protein distance of ankyrin1 and band4.1 in RBCs that had transited *Nc* ≈ 200 microchannels and static controls (Fig 5C). Our control measurements were in agreement with previous data^26^. Moreover, we observed a significant decrease in inter-protein spacing of ankyrin1 and band4.1 for *Nc* ≈ 200. Overall, these results suggest that vesiculation depletes specific proteins essential to RBC function, while most other proteins become slightly enriched as cells become denser. In particular, the network of bilayer-cytoskeleton linkers becomes tighter, consistent with the observed increase in effective bending modulus with *Nc* (see Fig 3D).

**Figure 5.**
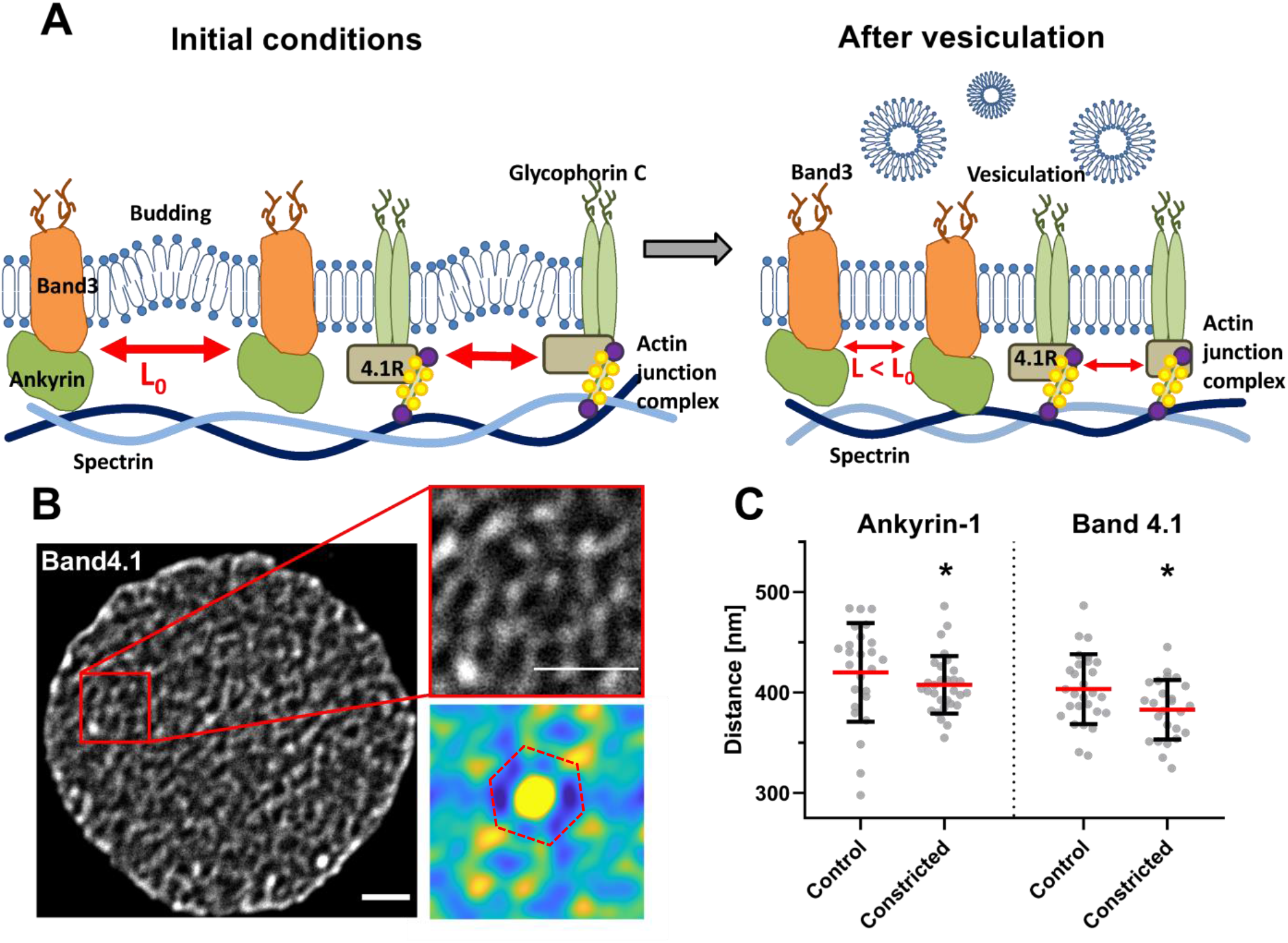
Passage through micro-constrictions increases the density of cytoskeleton-membrane anchoring complexes. **(A)**, A scheme explaining how vesiculation and membrane loss decreases the distance between the anchoring complexes that connect the membrane and the cytoskeleton. (**B, C**) STED microscopy was used to assess the distance between proteins that connect the membrane and the cytoskeleton (Ankyrin-1 and Band4.1) as indicated in Materials and Methods in static vs RBCs that crossed 200 microchannels (0.85µm width and 3µm length). In B, a representative image of a RBC labeled with an anti-Band4.1 antibody (scale bare 1 µm). The auto correlation matrix used to calculate the distance between two proteins is showed at the bottom. Red dashed line used to emphasize the hexagonal structure of the protein network observed in the autocorrelation image. Circles in C represent single cell measurements and bars indicate mean ± SD of at least 25 determinations. Statistically significant differences were measured using an unpaired t test with Welch’s correction and are indicated (*, p<0.05).

### Cytoskeletal integrity modulates RBC aging caused by microchannel crossing

To investigate the role of cytoskeletal integrity in mechanically induced RBC aging, we treated RBCs with drugs targeting different cytoskeleton components: Blebbistatin (Blebb) and Y27632 to inhibit myosin activity, and Latrunculin A (LatA) and Cytochalasin D (CytoD) to affect actin polymerization. These treatments increased cell deformability leading to shorter crossing times (Fig. 6A) and lower membrane bending modulus (Fig. 6B), with LatA producing the most significant effects. Besides, Y27632 and especially Blebb significantly reduced membrane tension (Suppl Fig. 5A). Given its more potent effects, we analyzed LatA treatment in more detail. As expected, LatA induced marked changes in RBC morphology as determined by electron microscopy (Suppl Fig 5B). Increasing LatA concentration accentuated both the decrease in RBC effective bending modulus (Fig 6B) and the number of released vesicles (Fig 6C). Consequently, cell volume loss with cumulative microchannel crossing was amplified by LatA treatment in a dose-dependent manner (Fig. 6D). Despite being initially more deformable and experiencing shorter crossing times than previously reported^27^, LatA-treated cells displayed increased crossing times (Fig 6E) and hemolysis (Fig 6F) for high *Nc*, likely due to LatA-induced, accelerated mechanical aging.

**Figure 6.**
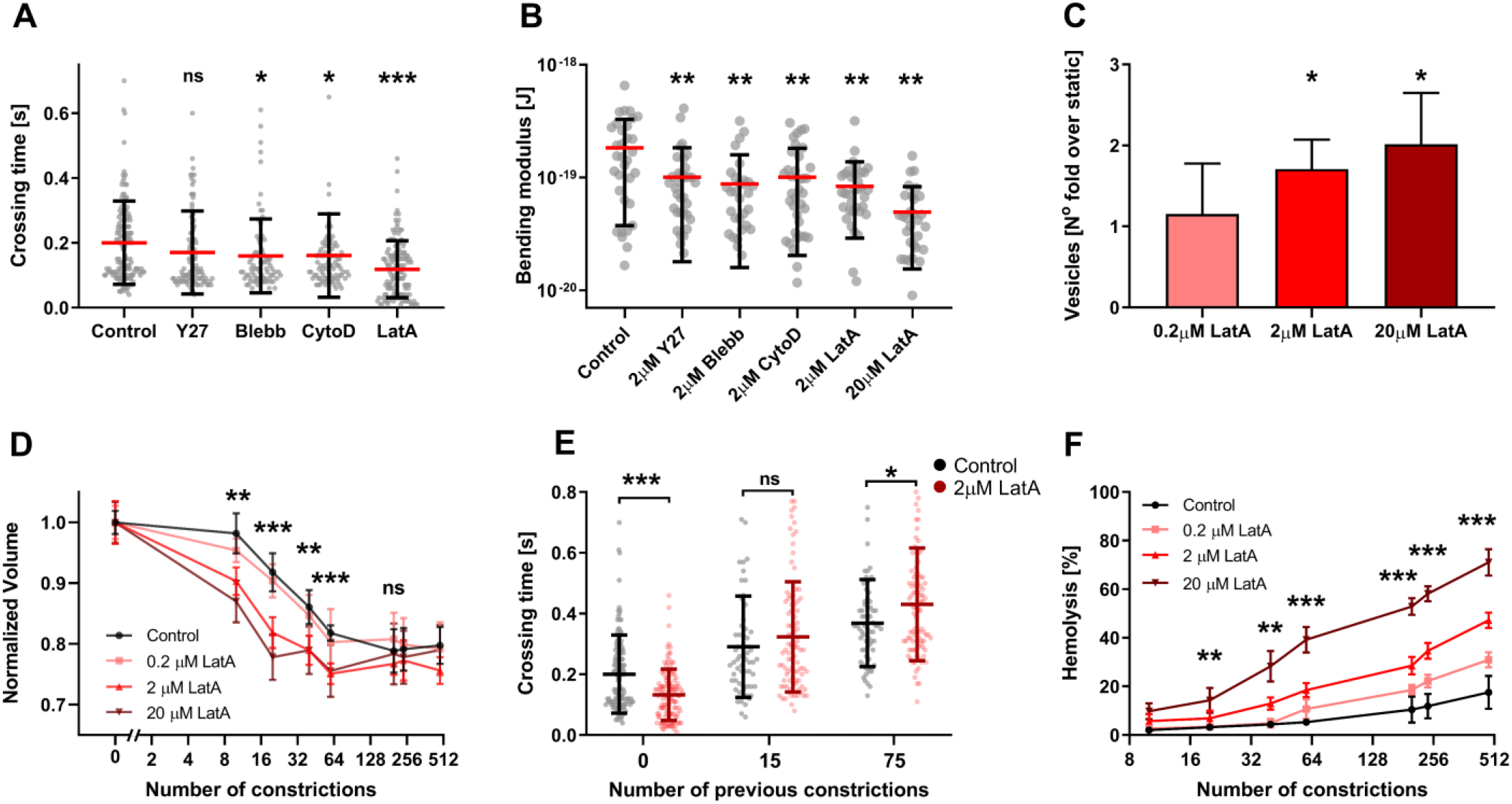
Cytoskeleton targeting drugs increase RBC deformability and demonstrate a relationship between vesiculation and cytoskeletal integrity. **A**, Microchannel (0.85µm width and 3µm length) crossing time in RBCs treated with Y27632, Blebbistatin, Cytochalasin D or Latrunculin A (in all cases 2 µM for 30 min), and control vehicle-treated cells. Circles represent single cell measurements and bars indicate mean ± SD of at least 25 determinations in 3 independent experiments. Statistically significant differences, as determined using an unpaired t test with Welch’s correction, are indicated (*, p< 0.05; ***, p< 0.001). **B**, ‘Effective’ bending elastic modulus of the cells treated in the same conditions as panel A or with 20 μM LatA. Circles represent single cell measurements and bars indicate mean ± SD of at least 10 determinations of 3 independent experiments. Statistically significant differences, as determined using a paired Wilcoxon test, are indicated (*, p< 0.05; **, p<0.01; ***, p< 0.001). **C**, Vesiculation caused by passing through a single micro-constriction by RBC treated with different LatA concentrations. Bars indicate mean ± SD of 4 independent experiments, p values were calculated using an unpaired t test with Welch’s correction and are indicated as (*, p< 0.05). **D**, Normalized cell volume vs. number of microchannels previously passed by RBCs after different doses of LatA treatment. Bars indicate mean ± SD of at least 20 determinations in 3 independent experiments. **E**, Micro-constriction crossing time after previous passage through 0, 15, or 75 microchannels in RBCs treated with 2 µM Latrunculin A and vehicle-control-treated RBCs. Circles represent single cell measurements and bars indicate mean ± SD of at least 20 determinations in 3 independent experiments. **F**, Hemolysis vs. number of microchannels previously passed by RBCs treated with different doses of LatA. Bars indicate mean ± SD of 6 independent experiments. For **D-F** statistically significant differences were measured between control and 2 µM LatA using an unpaired t test with Welch’s correction and are indicated (*, p< 0.05; **, p<0.01; ***, p< 0.001).

RBC oxidation is associated with changes in cytoskeletal integrity and cell deformability^28^. Accordingly, RBCs treated with H_2_O_2_ exhibited increased bending effective moduli and longer crossing times (Suppl. Fig 6), indicating a rigidized cytoskeleton. These results confirm that actin cytoskeleton integrity controls IES transit via RBC deformability. Moreover, impaired cytoskeletal integrity accelerates vesiculation, cell spherification, and other RBC aging hallmarks like hemolysis.

### RBC aging caused by microchannel crossing facilitates macrophage attack

To address whether mechanically induced RBC aging stimulates macrophages to target RBCs, we co-cultured THP-1-derived macrophages with RBCs after *Nc = 200* microchannel crossings. The RBCs were fluorescently labeled to tag macrophages that internalized RBCs (Fig 7A). This experiment revealed that the number of labeled THP-1 cells (Fig. 7B) and their average fluorescent intensity (Fig. 7C) increased significantly when they were co-cultured with mechanically-aged RBCs as compared to static RBCs (*Nc = 0*). Since RBC stiffness plays an important role in macrophage phagocytosis^29^, we treated RBCs with LatA or H_2_O_2_ to respectively promote or reduce their deformability. RBC treatment with H_2_O_2_ increased phagocytosis whereas LatA decreased it (Fig 7B-C).

**Figure 7.**
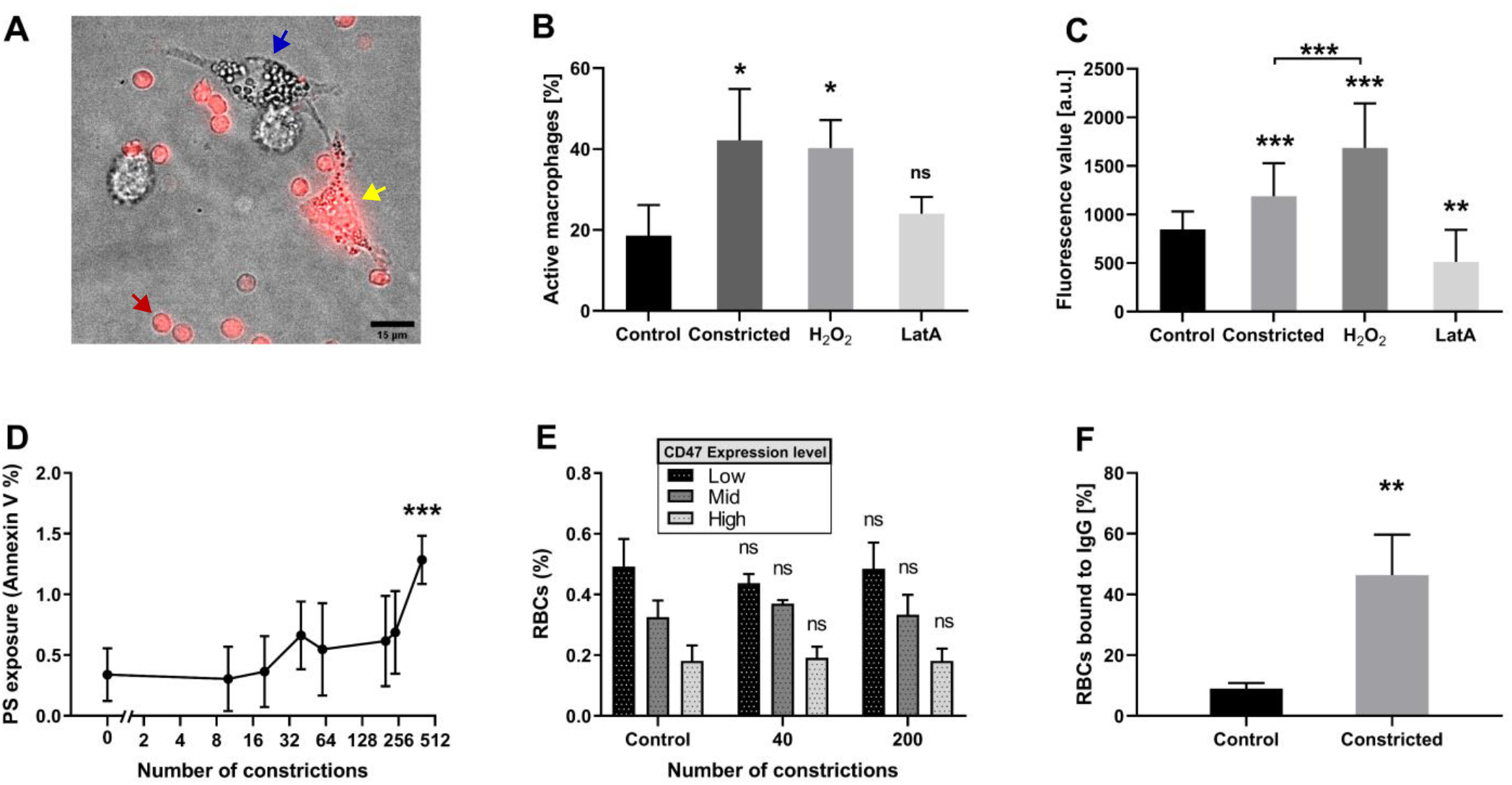
RBC passage through microchannels facilitates macrophage attack. **A**, THP-1 derived macrophages (blue arrow) were cultured with fluorescently-labeled RBCs (red arrow); an example of a macrophage that have incorporated fluorescence is shown (yellow arrow). **B**, Percentage of labelled macrophages detected after co-culture with fluorescent RBCs either static, after 200 microchannel crossings (0.85µm width and 3µm length) or treated with H_2_O_2_ (0.3 mM) or LatA (2 µM). **C**, Average fluorescence intensity of macrophages cultured with labelled RBCs treated as in panel B. In B and C, the bars represent mean ± SD of 4 independent experiments (*, p<0.05, using an unpaired t test with Welch’s correction). **D**, Exposure of phosphatidylserine (PS) on the outer plasma membrane of RBCs as determined by flow cytometry. A statistically significant difference was found between *Nc = 0* and *Nc = 480* as determined using an unpaired t test with Welch’s correction, and is indicated by (***, p< 0.001). Note also that the Y-axis only reaches 2%. **E**, Expression of transmembrane protein CD47 as determined by flow cytometry. No statistically significant differences were found as determined using an unpaired t test with Welch’s correction. **F**, The percentage of opsonized RBCs, static vs *Nc= 200*, that bound autoantibodies was assessed by flow cytometry. A statistically significant difference was found between *Nc = 0* and *Nc = 200* as determined using an unpaired t test with Welch’s correction, and is indicated by (**, p< 0.01). Means ± SD of 5 independent experiments are shown in **D**-**F**.

Next, we investigated whether cyclic microchannel crossing generated molecular “eat-me” signals in the RBC membrane. We measured phosphatidylserine (PS) exposure in the outer membrane leaflet^12^, finding a shallow increase in PS exposure that was statistically significant for *Nc = 512* (Fig 7D) but not for lower values of *Nc*. Thus, since we observed an increased RBC phagocytosis by macrophages for *Nc* = *200* (Fig 7B-C), PS exposure alone does not explain increased macrophage targeting of mechanically aged RBCs. We also measured the expression of the transmembrane protein CD47, whose decline with RBC age is associated with RBC clearance^13, 30^. However, we did not find any significant decrease in CD47 expression in RBCs that had crossed *Nc =* 40 or 200 microchannels as compared with static controls (Fig 7E), consistent with our proteomic data (see Fig 4).

Macrophage attack has also been associated with the clustering of specific membrane proteins like band3^31^, which could be active in mechanically aged RBCs based on our STED data (Fig 5). This targeting modality relies on the binding of autoimmune antibodies and their posterior recognition by macrophages via the Fc receptor^29, 32^. We opsonized static and *Nc = 200—*RBCs with plasma from the same blood sample, and measured the percentage of RBCs that bound immunoglobulin G (IgG). As shown in Fig 7F, mechanically aged RBCs exhibited a significantly higher level of IgG binding than controls. Thus, we concluded that mechanical RBC aging enhances macrophage attack by a combination of increased cellular rigidity and a higher association of autoantibody opsonization, with a possible contribution from increased PS exposure at later stages of the process.

## Discussion

This work demonstrates that the cumulative exposure of red blood cells (RBCs) to large-amplitude deformations promotes a decline in cell function similar to cell aging, and sheds light onto underlying mechanisms. Cyclic loading is known to lower RBC deformability^15, 16^, a mechanical function frequently used as an aging hallmark^18, 33^. We show that cyclic passage through IES (loading) also accentuates biological aging hallmarks like oxidation, ATP depletion, or hemolysis and, notably, degrades the cells’ oxygen-carrying efficiency. We investigate the molecular and structural origins of these phenomena and provide a nexus between cyclic loading and RBC targeting by macrophages, which is essential to remove aged RBCs from the circulation.

The largest deformations experienced by RBCs *in vivo* occur as they are filtered through the inter-endothelial slits (IES) of the spleen. Thus, we circulated RBCs through microfluidic devices that forced the cells to squeeze through narrow (< 1 μm) microchannels, causing large-amplitude deformations. The flow rate and microchannel dimensions were tuned to match *in vivo* measurements of IES morphology and crossing time^18, 20^. To extricate cyclic loading from other senescence-inducing processes like, *e*.*g*., continuous exposure to reactive oxygen species (ROS)^28^, we recirculated RBCs in our devices every 10 min, an interval 20 times shorter than the period between successive splenic passages *in vivo*. As RBCs were recirculated, their cumulative crossing of microchannels caused a cascade of biomechanical events that accelerated cell aging.

The earliest signs of RBC mechanical aging were detected after 15 load cycles and involved alterations in microchannel crossing dynamics (prolonged crossing time and more frequent cell folding vs. twisting motion). These changes were accompanied by a rise in cell oxidation and a decline in intracellular ATP. Concurrently, cell volume decreased, cell sphericity increased, and the bilayer-cytoskeleton complex stiffened, rendering the cells unable to transit the microchannels and causing hemolysis. The alterations became significant after 60 microchannel crossings, a comparable albeit smaller figure than the number of loading cycles that elicited mechanical RBC fatigue in previous studies^15, 16^. However, those studies had slower loading timescales (∼4 s ^15^ vs. 0.2 s) or wider microchannels (3 μm^16^ vs. < 1 μm) than splenic IES filtration^18, 20^, the process modeled in our microfluidic devices.

The observed changes were associated with the release of sub-micrometric vesicles. Vesiculation is observed in RBCs aging in vivo or stored for transfusion^34^. Mechanical deformation can trigger vesiculation by loading the linkages between the lipid bilayer and the cytoskeleton beyond their rupture strength. Simulations suggest these conditions are met at the trailing edge of RBCs passing through < 1-μm-wide microchannels^35^, and vesiculation has been reported in RBCs subjected to cyclic flow shear in vitro^36^. Besides, passage through constrictions elicits Ca^2+^ influxes^37^, an ion regulating RBC bilayer – cytoskeleton anchorage and implicated in vesiculation^38^. Our proteomics experiment indicates that calmodulin, a protein involved in Ca^2+^ regulation, is lost in mechanically aged RBCs, suggesting these cells have impaired Ca^2+^ activity, which could exacerbate vesiculation.

After 60 microchannel crossings, the RBCs stopped vesiculating and reached minimum volume and maximum sphericity, supporting the theory that vesiculation shuts off as the cells exhaust their available plasma membrane^36^. Our data suggest a mechanism for vesiculation tapering based on the density of bilayer-cytoskeleton linkages. We posit that the distance between linkages gradually decreases as vesicles are released, restricting bilayer fluctuations^39^. Consonantly, we observed that the distance between ankyrin1 and band4.1, two key bilayer-cytoskeleton crosslinkers, decreases significantly in mechanically aged RBCs. Moreover, the effective bending modulus of the bilayer-cytoskeleton complex increased with cyclic mechanical loading.

RBC aging and vesiculation have been linked to cellular oxidation, which is recognized to impair RBC deformability^40, 41^. Our data suggest that the interplays between these processes are more complex than previously appreciated because mechanical loading, in turn, also weakens the RBC antioxidant defenses. Circulating RBCs are constantly exposed to ROS released by exogenous (macrophages, endothelial cells) and endogenous sources. When not neutralized by the RBC antioxidant system, these ROS alter the RBC bilayer-cytoskeleton complex to increase cell stiffness. Our proteomic analysis revealed that mechanically aged RBCs lose a significant amount of antioxidant enzymes (e.g., catalase, peroxiredoxin-1, and peroxiredoxin-2). These results imply a novel positive feedback amplification loop between mechanical and oxidative damage.

Large-amplitude mechanical deformation promotes alterations that could favor the removal of RBCs from the circulation. First, as aged RBCs become less deformable, their microchannel crossing time increases significantly. *In vivo*, a prolonged IES crossing time would facilitate interactions between RBCs and macrophages in the spleen. Second, membrane composition changes in mechanically aged RBCs, discussed above, could modulate phagocytic signals recognized by macrophages. We found that THP-1-derived M1 macrophages displayed higher phagocytosis of mechanically aged RBCs compared to controls. Traditionally, PS^11, 12, 32^, CD47^13, 30^, and band3^10, 14, 32^ are considered as removal signals in senescent RBCs. However, our proteomics and flow cytometry measurements did not reveal marked changes in CD47 or band3 content in the membranes of RBCs that had squeezed through microchannels. Likewise, we only detected increased PS exposure after an extremely large number of deformation cycles. Thus, we concluded that cyclic loading may elicit non-traditional “eat-me” signals. Since target stiffness is known to affect phagocytosis efficiency^29^, we hypothesized that increased oxidation of membrane proteins in mechanically aged RBCs and its associated cell stiffening facilitate macrophage attack. Consistent with this hypothesis, we observed that H_2_O_2_ treatment increases RBCs stiffness and promotes phagocytosis. In addition, mechanically aged RBCs had higher opsonization efficiency, which can contribute to RBC phagocytosis via the Fc receptor in macrophages. Autoantibody binding has been related to band3 oxidation^32^. Therefore, we surmise that the increased phagocytosis of mechanically aged RBCs arises from the combination of auto-antibody opsonization and increased cell rigidity and oxidation.

Vesiculation allows RBCs to shed membrane fragments containing pro-phagocytic molecules like PS or denaturalized band3^42^. In this regard, vesiculation is believed to protect RBCs until it progressively shuts off with cell age. We tested whether treating RBCs with LatA, a drug that drives F-actin depolymerization, would prolong vesiculation, preserve the cells’ deformability, and prevent the aging effects induced by cyclic loading. However, while it initially increased cell deformability and protected the cells from macrophage attacks, LatA treatment also accelerated vesiculation shut off and the adoption of a fragile spherical phenotype. Overall, our results imply that vesiculation introduces a clock in RBC functional decline, winding up with impaired cell deformability, reduced antioxidant protection, decreased gas transport and exchange, and increased macrophage phagocytosis. Notably, we have shown that this clock can be hastened by cyclic mechanical loading.

## Materials and Methods

### RBC isolation

Human blood from healthy donors was obtained in Heparin treated BD Vacutainer tubes from the San Diego Blood Bank. Upon reception, red blood cells (RBCs) were isolated using a density gradient technique^43^. First, a 15 ml centrifuge tube (Falcon tube, Fisher Scientific) was filled with 3 ml of Histopaque 1119 (Sigma Aldrich); then, a 3 ml layer of Histopaque 1077 (Sigma Aldrich) was added on top and finally 6 ml of whole blood were carefully added to prevent the three layers from mixing. The tube was centrifuged at 700 g for 30 min at room temperature; the top layers containing plasma, mononuclear cells, and granulocytes were discarded and the RBCs were resuspended in PBS (Dulbecco’s Phosphate Buffered Saline, Sigma Aldrich). The tube was centrifuged again at 200 g for 10 min at room temperature and the supernatant cells were removed to discard platelets and damaged RBCs. Finally, RBCs were resuspended in AS-3 media^44^ (dextrose, sodium chloride, adenine, citric acid, sodium citrate and sodium phosphate were all bought from FisherScientific) in a 5% hematocrit concentration and stored at 4° C until used. All the experiments were conducted within one week of RBC isolation. Separation of denser and lighter RBCs was performed by discontinuous gradient of Histopaque. A layer of RBCs was carefully added on top and centrifuged for 15 minutes at 700 g. Layers were aspirated carefully and only bottom(denser) and top (lighter) cells were kept.

### Microfluidic device fabrication

A series of polydimethylsiloxane (PDMS) microfluidic devices with multiple constrictions of customizable geometry were designed and manufactured in the San Diego Nanotechnology Facility of UC San Diego. A two-step photolithography process combining both positive and negative photoresist was used to fabricate the mold of the device, resulting in a two-layer device. The thicker layer, used for the inlet and outlet regions, was made with negative photoresist to allow for minimized flow resistance and RBC deformation outside of the constrictions. The central part of the device containing the constrictions required higher resolution which was obtained leveraging the thinner layer of positive photoresist.

The process started by cleaning a 4-inch silicon wafer (University Wafer, South Boston, MA) in a sonic bath of acetone for 5 min; then it was rinsed in methanol, isopropanol and finally water. Subsequently, the wafer was dehydrated at 180° C for 10 min and its surface activated using a plasma treatment (PVA TePla PS100) for 5 min. A coat of MCC Primer 80/20 (Microchemicals, Westborough, MA) was applied to improve the attachment of the photoresist to the wafer. After a short (2 min) bake at 110° C, the first layer of SU8 2050 negative photoresist (Microchemicals, Westborough, MA) was applied by spin coating at 3000 rpm for 30 sec. After 6 min of soft bake at 95°C, the inlet and outlet regions of the device were exposed using a MLA150 (Heidelberg Instruments Mikrotechnik GmbH, Germany) under a laser with 375 nm wavelength and a dose of 3000 mJ/cm^2^. The post exposure bake was carried at 95° C for 5 min and then the wafer was developed using SU8 developer (Microchemicals, Westborough, MA). The second part of the process started by applying again a coat of MCC Primer and then spin coating AZ12XT-20PL-05 (Microchemicals, Westborough, MA) for 30 sec at 800 rpm. After soft baking the sample for 2 min at 110° C, the whole device, including inlet/outlet and central region was exposed using a dose of 200 mJ/cm^2^. Afterwards, the post exposure bake was carried on for 1 min at 90° C and the wafer was developed using AZ300 MIF (Microchemicals, Westborough, MA). The heights of the channel were verified using a Dektak 150 surface profilometer (Veeco Instruments Inc. Plainview, NY) and the wafer was passivated with tridecafluoro-1,1,2,2-tetra-hydrooctyl-1–trichlorosilane e (Gelest, Morrisville, PA) for 15 min inside a vacuum chamber to prevent PDMS adhesion to the wafer.

PDMS replicas of the device were made by casting a previously degassed mixture of the PDMS oligomer and crosslinking agent (Sylgard^®^184, Dow Corning Inc, Midland, MI) in a 10:1 (w/w) proportion on the passivated silicon wafer. The sample was then cured at 65° C overnight. The next day the master was peeled off from the wafer, cut into several single devices and the inlet and outlet holes were punched (2.5 mm) with a biopsy puncher (Miltex, Integra Lifesciences, Plainsboro Township, NJ). Finally, we activated the surface of both coverslip (Corning, 24×60 mm and thickness 1.5 mm) and PDMS chip under a UV ozone lamp (Model 30, Jelight Co. Irvine, CA) for 4 min with an oxygen inflow of 0.2 l/min and bonded them together at 65° C for a minimum of 4 h before they were ready to be used.

### RBC morphological determinations

To calculate the volume and the surface of RBCs we used a method based on Fluorescence exclusion as previously described^45, 46^. Briefly, RBCs were suspended at a concentration of 10^4^ cells/ml in the AS-3 media supplemented with 1 mg/ml of FITC-Dextran (Sigma-Aldrich). The solution containing the cells was introduced into one of the microfluidic devices described previously and imaged using a Leica DMI 6000B inverted phase contrast microscope (Leica Camera, Wetzlar, Germany). Chambers’ heights [4.5-5 µm] were rigorously characterized using profilometer (Dektak 150 Surface Profiler, Veeco). In the fluorescent field, RBCs and PDMS pillars appeared dark whereas the rest of the chamber, where the Dextran was present, resulted in brighter regions. Image analysis was performed using a in house MATLAB (The Math Works Inc., Natick, MA, USA) scripts. For each image, a calibration using PDMS pillars was used to obtain the relationship between fluorescence intensity and chamber height as described previously^45^. The linear relationship between the height of the object and the intensity is described by: *IB = α · h + IP*, where IB and IP are the background and PDMS pillar fluorescence respectively and *h* the height of the chamber. To measure single cell volume, a region of interest was automatically defined around each cell where the background was calculated; then *α* was obtained and finally the heights of the pixels conforming the cell were obtained. To calculate the surface, an equatorial symmetry for the cells was assumed, and an enclosed area was generated using half of the pixel heights. The cell was considered valid if the difference between the volume computed from the enclosed area and the volume calculated from the pixel height was smaller than a threshold (10%). Echinocytes were discarded from surface calculation due to the limitations of the equatorial symmetry assumption. Our *V* and *S* measurements produced results in agreement with previously obtained data by other techniques^47, 48^ (Suppl Fig. 2A-B).

### Determination of RBCs ‘effective’ bending modulus and membrane tension

To measure the mechanical properties of RBCs we used thermal flickering spectrometry as described in ^23^. Using this technique we captured the contributions from both the lipid bilayer and the cytoskeleton. RBCs were diluted into culture medium at 10^4^ cells/ml and seeded into a thin coverslip (Corning). Leica DMI 6000B inverted phase contrast microscope (Leica Camera, Wetzlar,Germany) controlled by a dedicated workstation connected to a Zyla3-Tap Enclosed C-mount 16 bit camera (Andor Technology, Belfast, UK) was used for high speed (300 fps) image acquisition. Imaging was performed in a controlled environment at 37 °C by using the setup described above. A MATLAB (The Math Works Inc., Natick, MA, USA) code was written to analyze the images and obtain the mechanical properties of the cells. First, the RBC contour at the cell equator was detected for each frame using the method described in ^49^. Then, the power spectrum of mean square mode amplitudes was obtained by applying the Fourier transform and, from these data, the ‘effective’ bending modulus (κ) and tension (σ) were fitted using the following equation:

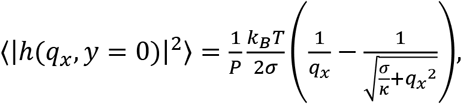

where k_B_ is the Boltzmann constant, *T* is the absolute temperature, and *P* is the mean perimeter of the RBC contour. This equation was derived from the energy of deforming a flat sheet^50^ and describes shape fluctuations of the cell’s equator in only a limited range of modes. On one hand, smaller modes are excluded because they are affected by the geometry of the surface and an important deviation from the spherical harmonics expression. On the other hand, high modes, with smaller wavelengths, are affected by noise and their fluctuations lie outside the spatial and temporal resolution of the experiment. For this reason, we use modes 6 to 18 for fitting the equation above.

### Flow Cytometry

Flow Cytometry was used to measure the vesicles released by RBCs. Briefly, after finishing every experiment, RBC samples were resuspended at 10^6^ cells/ml in PBS and analyzed using an Accury C6 (BD Biosciences, USA). Blank samples, containing only PBS, were used to set the forward and side scattering thresholds to limit noise measurements, that would reflect in an incorrect reading in the number of vesicles. Manual gating for the RBCs and samples was performed using a user made MATLAB (The Math Works Inc., Natick, MA, USA) code.

Phosphatildylserine exposure on the outer leaflet of RBCs was measured using the Annexin V Alexa Fluor™ kit (V13241, ThermoFIsher). RBCs were resuspended at a concentration of 10^6^ cells/ml in the Annexin V binding buffer. Then, 5 µL of Alexa Fluor^®^ 488 annexin V and 1 µL of Propidium Iodide were added into 100 µL of cell suspension and incubated for 30 min. Finally, cells were centrifuged and resuspended in the binding buffer for analysis using the flow cytometer. Negative controls were used to set up the threshold for the detection of positive cells.

To measure the expression levels of CD47 we incubated the cells with anti-CD47 (Clone B6H12, BDbiosciences) and 2% BSA for 1 h at room temperature. We then washed thrice in PBS and incubated the cells with the secondary antibody (Goat anti-Mouse IgG Alexa Fluor 488, ThermoFisher). Shortly after, the samples were analyzed using the flow cytometer. Appropriate negative controls were prepared to differentiate positive cells.

To measure the percentage of opsonized RBCs we first treated cells with 2% BSA for 1 h, washed with PBS multiple times and incubated with plasma from the same blood sample for 1 h. Finally, RBCs were incubated with anti-Human IgG Alexa Fluor 488 (Thermo Fisher) for 2 h, fixed with 4% PFA and analyzed using the flow cytometer. Negative controls were used to establish a fluorescent threshold.

### Stimulated emission depletion (STED) microscopy

RBCs samples were bonded to Nº 1.5 glass coverslips treated with 100 ng/ml Poly-L-Lysine (Sigma-Aldrich) for 1 h. Cells were fixed using 4% PFA for 15 min, washed multiple times in PBS and permeabilized with 0.1% Triton-X (Sigma-Aldrich) for 10 min. Non-specific antibody binding was prevented by incubating with 2% BSA (Thermo Fisher) for 2 h. Cells were incubated with the respective primary antibody (rabbit anti human, anti-ANK1 or anti-EPB41, Sigma-Aldrich) overnight. Finally, a secondary antibody (goat anti-rabbit IgG, Thermo Fisher) was added for 1h and samples were fixed using ProLong Diamond (Thermo Fisher) for at least 24 h. Samples were imaged in the Microscope Core Facility at UCSD (NS047101) using a Leica SP8 confocal with STED and lightning deconvolution. Post processing of the images and data analysis was performed with an in-house designed MATLAB script. Briefly, the autocorrelation of the images was obtained and the distance between the two higher peaks was used to calculate the distance in between protein complexes.

### Hemolysis

The content of hemoglobin in the supernatant was used to measure the percentage of RBCs lysis. Hemoglobin concentration was measured analyzing absorbance at 415 nm in a UV-Visible spectrophotometer (Biomate 3s, ThermoFisher). Before starting the different experiments, a sample containing the same concentration of RBCs was used to later calibrate the absolute hemolysis lysing the cells with 1% SDS. After finishing every experiment RBCs were pelleted down by centrifuging at 1000 rpm for 2 min (Eppendorf, 5415C) and the supernatant absorbance was assessed and normalized using the maximum hemolysis value.

### RBC oxidation

RBCs oxidation was assessed by measuring the intracellular content of Methemoglobin (MetHb) as previously described^51^. RBCs obtained from the different samples were washed multiple times with PBS to remove any extracellular hemoglobin from lysed cells. Then, RBCs were resuspended in PBS at a final concentration of 10^7^ cells/ml, lysed with 1% SDS and the absorbance at 645 nm was measured using a UV-Visible spectrophotometer (Biomate 3s, ThermoFisher). The solution was then treated with potassium hexacyanoferrate (K_3_[Fe(CN)_6_], ThermoFisher) to completely oxidized Hb into MetHb and the absorbance was measured again. Finally, the percentage of MetHb was obtained by normalizing the two measurements at 645 nm.

### ATP concentration

The intracellular ATP concentration was measured using the Luciferin technique (ATP Determination Kit; ThermoFisher). Briefly, 10^5^ cells were extracted from each sample and washed with PBS multiple times to remove any cell remaining in the supernatant. The standard reaction solution containing luciferase and luciferin was prepared according to the manufacturer guidelines and introduced in a black with clear bottom 96-well plate (Corning). To prevent any possible ATP degradation, RBCs samples were lysed with DI water and quickly added to each well. When ATP reacted with the standard solution, light was emitted with a wavelength of 560 nm and captured with a luminometer (Tecan infinite M1000 Pro). A standard reference curve was made from different ATP concentration from an ATP vial provided with the kit. This curve was used to measure the ATP concentration for each sample. Finally, to obtain the intracellular ATP concentration, the average measured concentration was normalized by the total number of cells lysed and the mean volume obtained in previous experiments. ATP measured concentrations were similar to previous studies^52^.

### RBCs recirculation and crossing time determination

Flow was driven through the device at a fix flow rate (100µl/h) using a peristaltic pump (IPC, ISMATEC) (Suppl Fig 1F); wider channels (5 μm) were added at both sides of the microchannel array to ensure that microchannel clogging by stuck RBCs did not significantly affect the pressure difference across the array. The maximum number of constrictions crossed by the RBCs was estimated based on the flowrate and the volume of fluid inside the circuit. Therefore, *Nc* represents the maximum number of constrictions that RBCs could have undergone for a period of time. RBCs were recirculated at room temperature.

To measure the crossing time RBCs were washed multiple times in PBS and resuspended at 1% HC in AS-3. The device was left running for 15 min before the image acquisition started to allow the flow to stabilize. The passage of RBCs through misconstructions was imaged using a DMI 6000B inverted phase contrast microscope (Leica Camera, Wetzlar, Germany) equipped with ahigh frame rate (100 fps) camera (Zyla3-Tap Enclosed C-mount 16 bit, Andor Technology, Belfast, UK). The crossing time was manually calculated as the difference between the first and last frame where the RBCs contacted the constriction.

### Proteomics analysis

Control RBCs and cells transited through 200 constrictions were collected and washed multiple times with PBS. Then, cells were lysed with DI water and centrifuged at 14000 g for 10 min to collect the cell membranes. The pellet was resuspended in DI water and washed multiple times to remove as much Hemoglobin as possible. The total protein concentration was measured using a Micro BCA Protein Assay Kit (Thermo Fisher) and resuspended at a final concentration of 120 µg/ml for all samples. Samples were submitted to the Biomolecular and Proteomics Mass Spectrometry Facility (BPMSF, S10 OD016234) at UCSD and analyzed using a label free quantification. The Benjamini-Hochberg correction method^53^ was used to adjust the p-values. Protein set enrichment analysis was performed using Metascape on the KEGG, Canonical pathways, Gene Ontology, Reactome, and CORUM databases^25^.

### Phagocytosis assay

Macrophages were differentiated from THP-1 monocytic cells based on^54^. Briefly, 5×10^5^ THP-1 cells/ml were treated with 25 ng/ml phorbol myristate acetate (PMA, ThermoFisher) in RPMI 1640 with 10% FBS. After 72 h, the media was renewed and a rest period of 72 h was performed before cytokine activation. To polarize macrophages into M1 phenotype, cells were treated with 50 ng/ml of IFNγ (eBiosciences) and 200 ng/ml of LPS (eBiosciences) for 72 h. At this point cells were already activated and ready to interact with RBCs.

RBCs from control or cells transited through 200 constrictions were collected washed three times with PBS and resuspended at a concentration of 10^6^ cells/ml. Following, RBCs were stained with CellTracker Deep Red CMFDA Dye (Invitrogen) for 30 min at 37ºC and washed multiples times with PBS. Finally, RBCs were opsonized with plasma from the same blood samples for 30 min, cocultured with macrophages in RPMI +10%FBS for 48 h and fixed afterwards for imaging using 4% paraformaldehyde (PFA, ThermoFisher). Cells were imaged using a DMI 6000B inverted phase contrast microscope (Leica Camera, Wetzlar, Germany) and active macrophages were assessed by measuring its fluorescence. To measure the mean fluorescence of each macrophage, cells were manually gated and the mean intensity of inner pixels was calculated.

## Acknowledgements

We thank to the members of the EMT and Tumor Invasion Group from the IMIM at Barcelona for their advice and help. This work was performed in part at the San Diego Nanotechnology Infrastructure (SDNI) of UCSD, a member of the National Nanotechnology Coordinated Infrastructure supported by the National Science Foundation (Grant ECCS-1542148). This work was funded by Grant NSF CBET - 1706436/1948347 and NSF CBET - 1706571. Yi-Ting Yeh would like to acknowledge American Heart Association (18CDA34110462).

