## Supplementary material for "Cyclic mechanical stresses alter erythrocyte membrane composition and microstructure and trigger macrophage phagocytosis": Supplementary Information.pdf

**Analysis of the role of mechanical stress on red blood cell aging**

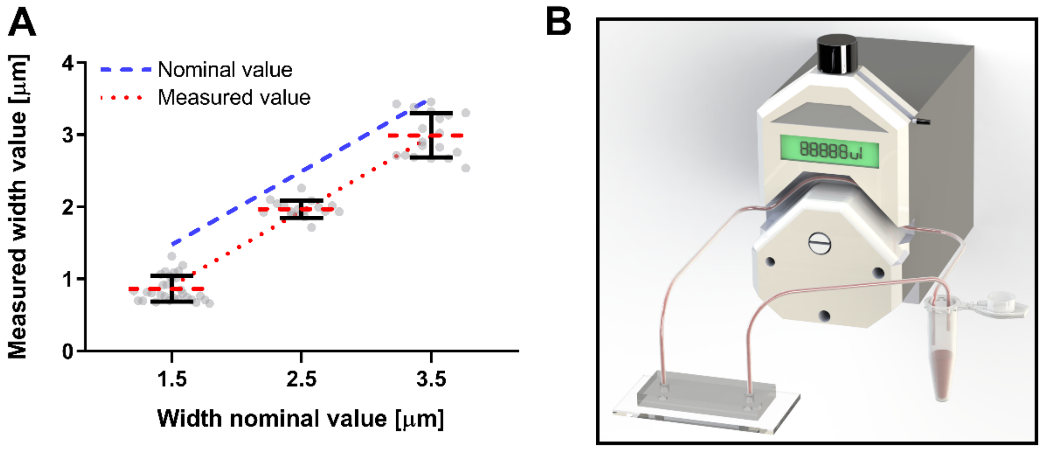


**Supplementary Figure 1. Characterization of the device**. **A**, Device's geometry was determined using scanning electron microscopy. Measured widths were 0.5 µm smaller than the nominal design value. **B**, Set up used to recirculate cells through the device multiple times using a peristaltic pump.

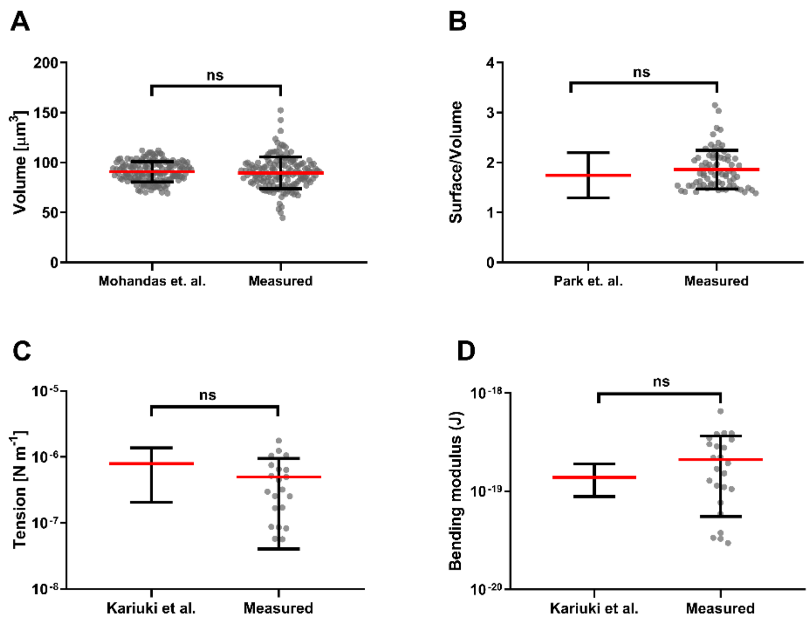


**Supplementary Figure 2. Comparison of measured cellular parameters with previous determinations.** Single cell volume (**A**) and surface to volume ratio (**B**) were measured using fluorescence exclusion and compared with previous studies[2, 3]. Membrane flickering spectrometry was used to quantify membrane tension (**C**) and bending modulus (**D**); measurements also agreed with previous studies[4]. Statistically non-significant differences were observed using an unpaired t test with Welch's correction.

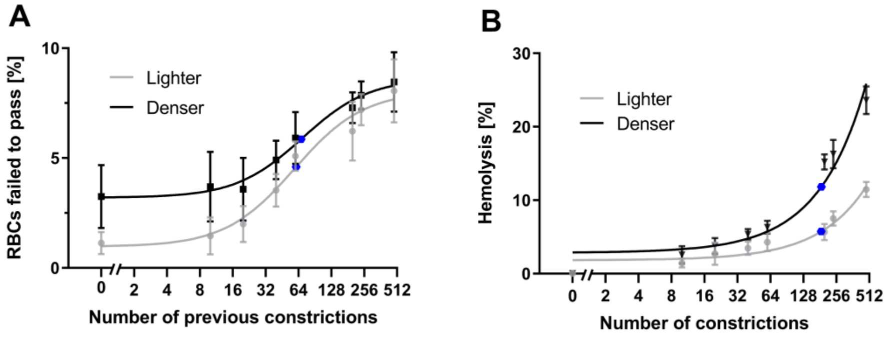


**Supplementary Figure 3. Repetitive passage increased the failure of RBCs to cross the microchannels**. The figures show the retention rate (**A**) and hemolysis (**B**) for both denser and lighter RBC subpopulations with respect to previous or total passages. Error bars represent mean ± SD for at least 20 cells of 3 independent experiments. Data was modeled using the Hill-dose-response equation (A) and a double exponential (B). The blue dot corresponds to the point with 50% change.

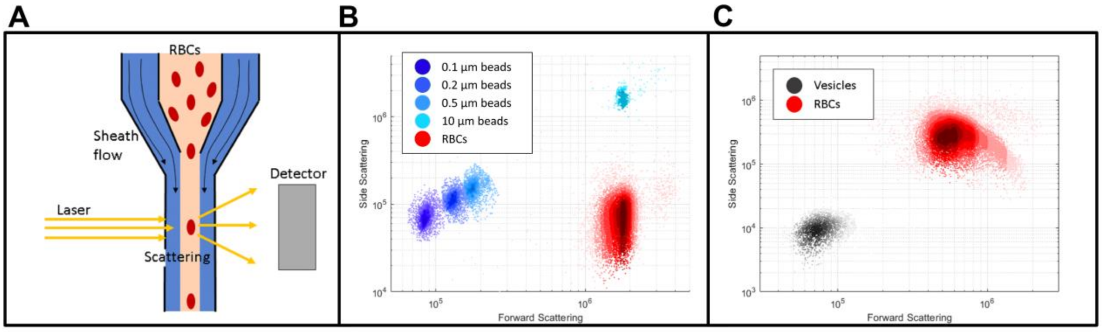


**Supplementary Figure 4. Determination of RBCs vesiculation**. **A**, Flow cytometry was used to quantify vesiculation. Forward and side scattering were used to differentiate the vesicles from the cells. The set up was calibrated using beads of know sizes (0.1-10 µm) and compared with RBCs (**B**). Samples collected from the device were analyzed and a population of vesicles, comparable to beads between 0.1 -0.5 µm in diameter, was obtained (**C**).

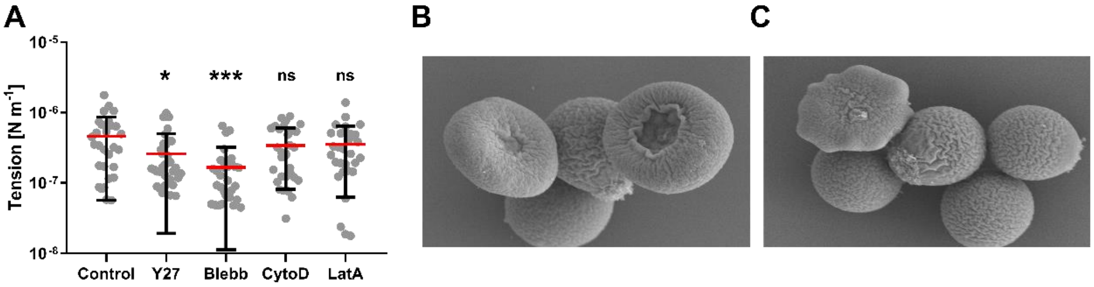


**Supplementary Figure 5. Effect of cytoskeleton inhibitors on RBCs**. **A**, Membrane flickering spectrometry was used to measure the tension of RBCs treated for 30 min with Y27632, Blebbistatin, Cytochalasin D and Latrunculin A (all 2 µM). Single cell measurements are represented as circles; error bars represent mean ± SD. Statistically significant differences, as determined using a paired Wilcoxon test, are indicated (*, $p < 0.05$; ***, $p < 0.001$). Pictographs taken with scanning electron microscopy of control (**B**) and LatA-treated (**C**) RBCs.

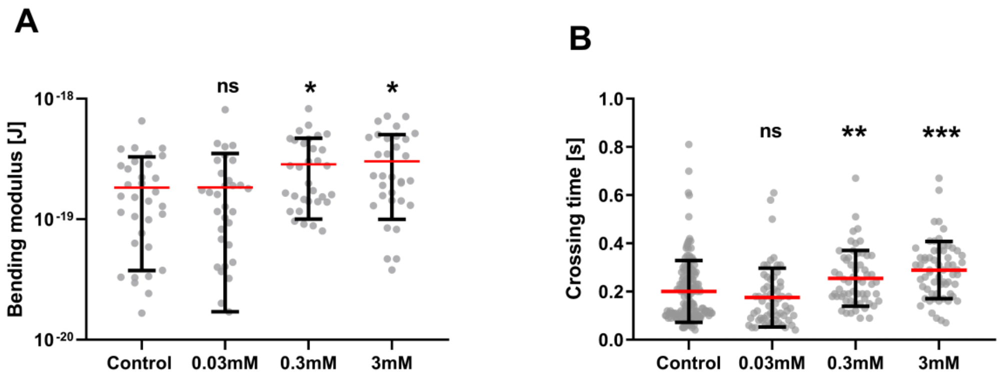


**Supplementary Figure 6. Oxidation modifies RBC physical properties. A,** 'Effective' bending modulus measured with membrane flickering spectrometry of cells treated for 30 min with different concentration of $H_2O_2$. **B,** RBCs crossing time was assessed for different concentrations of $H_2O_2$. Statistically significant differences determined using an unpaired t test with Welch's correction, are indicated (*, $p< 0.05$; **, $p< 0.01$; ***, $p< 0.001$). Single cell measurements are represented as circles; error bars represent mean ± SD.
